# Molecular basis for the activation of *Pseudomonas aeruginosa* MsbA by Zn^2+^

**DOI:** 10.1101/2024.12.31.630943

**Authors:** Jixing Lyu, Hanieh Bahramimoghaddam, Tianqi Zhang, Gaya P. Yadav, Minglei Zhao, David Russell, Arthur Laganowsky

## Abstract

Proteins involved in the biogenesis of lipopolysaccharide (LPS), a lipid exclusive to Gram-negative bacteria, are promising candidates for drug discovery. Specifically, the ABC transporter MsbA plays a crucial role in translocating an LPS precursor from the cytoplasmic to periplasmic facing leaflet of the inner membrane, and small molecules that inhibit its function exhibit bactericidal activity. Here, we use native mass spectrometry (MS) to determine lipid binding affinities of MsbA from *P. aeruginosa* (PaMsbA), a Gram-negative bacteria associated with hospital-acquired infections, in different conformations. We show the ATPase activity of the transporter is stimulated by Zn^2+^ and successfully trapping the protein with vanadate requires Zn^2+^ not Mg^2+^, which is necessary to trap MsbA from *E. coli*. We also present cryogenic-electron microscopy structures of PaMsbA in occluded and open outward-facing conformations determined to a resolution of 2.98 and 2.72 angstroms, respectively. The structures reveal a triad of histidine residues and mutation of these residues abolishes Zn^2+^ stimulation of PaMsbA activity. Together our studies provide detailed insight into PaMsbA structure, lipid binding preferences, and uncover a mechanism through which Zn^2+^ promotes the dimerization of the transporter, resulting in enhanced ATPase activity.

## Introduction

The increasing prevalence of antimicrobial-resistant pathogens, which are often associated with nosocomial infections, imposes a substantial strain on healthcare systems.^1^ A group of bacteria have emerged that evade the lethal effects of antibiotics, commonly referred to as “the ESKAPE bugs.”^2,3^ ESKAPE is an acronym for *Enterococcus faecium*, *Staphylococcus aureus*, *Klebsiella pneumoniae*, *Acinetobacter baumannii*, *Pseudomonas aeruginosa*, and *Enterobacter species*. Specifically, *P. aeruginosa* ranks among the top three leading causes of opportunistic human infections, contributing to 10-20% of hospital-acquired infections.^4,5^ Metals play key roles during infection and zinc has been shown to play important roles for *P. aeruginosa*, including virulence, host organism colonization, and antibiotic resistance.^6,7^ Following invasion by infecting bacteria, the body promptly triggers a nutritional immune response, leading to a decrease in free zinc levels in the blood and tissues.^6,8^

A number of proteins involved in LPS synthesis and transport in Gram-negative pathogens represent attractive antibacterial targets.^9,10^ In particular, MsbA is gaining recognition as a promising antibiotic target because of its vital involvement in the biogenesis of LPS, a cell component critical for bacterial survival.^11–13^ Deletion of the *msbA* gene is fatal for bacteria, underscoring the significance of targeting this transporter.^14,15^ Several groups have developed small molecule inhibitors of MsbA that display bactericidal activity.^16–20^ However, the role of lipids in modulating selective interaction of inhibitors with MsbA is not well understood.

MsbA is an essential ATP-binding cassette transporter in gram-negative bacteria that plays an important role in lipopolysaccharide (LPS) biogenesis.^21,22^ MsbA functions as a homodimer, and each subunit contains a nucleotide binding domain (NBD) and transmembrane domain (TMD), composed of six transmembrane helices per subunit.^16,23–34^ The transporter uses the hydrolysis of ATP, which binds to the NBD, to power a conformational change from an open, inward-facing conformation to an outward-facing conformation.^29^ This conformational change is a critical step in the transport of LPS, flipping the lipid from the cytoplasmic leaflet to the periplasmic leaflet of the inner membrane. While MsbA from *E. coli* (EcMsbA) has been extensively studied,^16,23–34^ the transporter from *P. aeruginosa* (PaMsbA), sharing 40% sequence identity, is relatively less explored,^15^ particularly in terms of its structure and lipid binding preferences. Here, we use native mass spectrometry to characterize PaMsbA-lipid interactions and identify conditions to trap the transporter, revealing an unexpected role of zinc as an important metal cofactor. We then determine cryoEM structures of PaMsbA in two different conformations leading to the identification of a triad of histidine residues. Biochemical studies show these residues are important for stimulation of the transporter by zinc, which is likely the result of promoting dimerization of the NBDs.

## Results

### Determination of PaMsbA-lipid binding affinities

To better understand the lipid binding preferences of PaMsbA, we first prepared the transporter for native mass spectrometry (MS) studies. In our previous study,^34^ we optimized the purification of EcMsbA, allowing us to detect the binding of small molecules, like copper ions (65 Da), to the transporter. Following these procedures, a well resolved mass spectrum of homodimeric PaMsbA in the C_10_E_5_ detergent was obtained with a measured mass (133.15 kDa) in agreement with the theoretical mass (133.15 kDa) (**Figure 1**). Unlike EcMsbA, no adducts consistent with the mass of copper ions bound to the transporter were observed. This result is anticipated as the N-terminal sequence of PaMsbA differs from EcMsbA, wherein the N-terminal sequence (MHD…) of EcMsbA binds specifically to Cu^2+^.^34^

**Figure 1.**
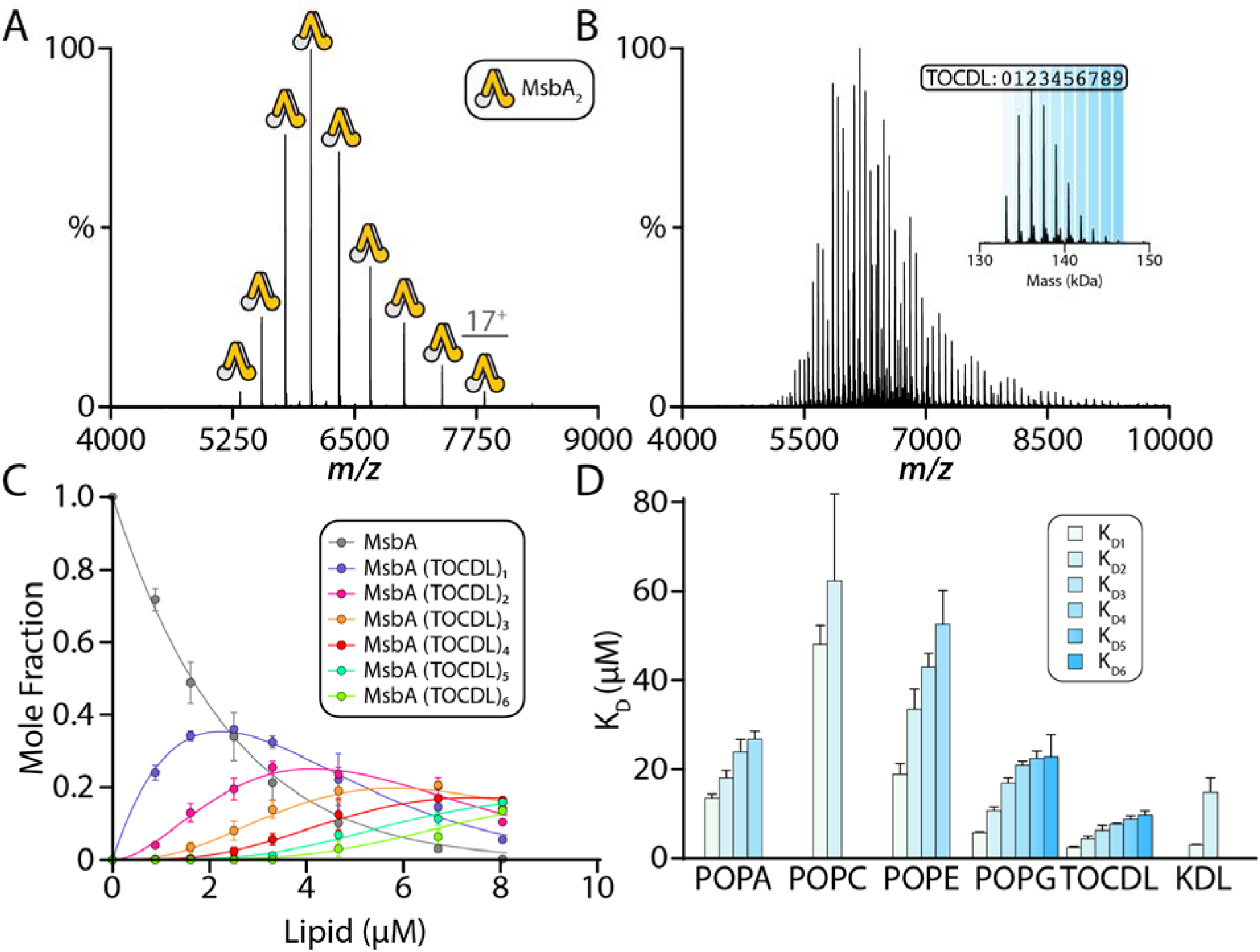
Determination of the lipid binding affinities of PaMsbA. A) Native mass spectrum of PaMsbA in the C_10_E_5_ detergent. B) Native mass spectrum of 4.4 µM PaMsbA with 6 µM TOCDL. The deconvoluted mass spectrum is shown in the inset. C) Plot of the mole fraction of PaMsbA(TOCDL)_0-6_ determined from a titration series (dots) and resulting fit of a sequential ligand binding model (solid lines). D) Equilibrium dissociation constants (K_D_s) for different lipids binding to PaMsbA. Reported are mean and standard deviation (*n* = 3).

We next characterized the binding of lipids to PaMsbA. For these studies, we focused on the major lipids found in *Pseudomonas*^35^ and selected 1,1⍰,2,2⍰-tetraoleoyl-cardiolipin (TOCDL), and phosphatidic acid (PA), phosphatidylcholine (PC), phosphatidylethanolamine (PE), phosphatidylglycerol (PG), and phosphatidylserine (PS) containing 1-palmitoyl-2-oleoyl (PO, 16:0-18:1) acyl chains. We also included 3-deoxy-D-*manno*-oct-2-ulosonic acid (Kdo)_2_-lipid A (KDL), an LPS precursor known to stimulate the ATPase activity of MsbA.^15,36–38^ The mass spectrum of PaMsbA in the presence of ∼1.5 molar equivalents of TOCDL showed a number of lipids bound to the transporter. More specifically, up to nine TOCDLs bound to PaMsbA (**Figure 1B**). PaMsbA was titrated with TOCDL and the resulting mass spectra were processed to determine the equilibrium dissociation constants (K_D_s) for PaMsbA binding one to six TOCDLs (**Figure 1C-D**). Titration with other lipids was performed in an analogous fashion, showing that TOCDL (K_D1_=2.4 ± 0.2 μM) and KDL (K_D1_=3.0 ± 0.2 μM) bind to PaMsbA with a relatively high binding affinity compared to other lipids in this study (**Figure 1D and Supplementary Table 1**). However, the TOCDL and KDL binding affinity of PaMsbA is weaker compared to EcMsbA with a K_D1_ of 1.6 and 0.6 μM, respectively.

### Discovery of zinc as an important PaMsbA cofactor

Following the determination of lipid binding affinity in the absence of any nucleotide, our goal was to investigate how lipid binding affinity in PaMsbA is influenced by its conformation. We first attempted to trap PaMsbA using adenosine 5’-triphosphate (ATP) and vanadate in the presence of Mg^2+^, a procedure we used to successfully trap EcMsbA in an outward facing conformation (**Figure 2A, top panels**).^34^ Interestingly, PaMsbA remained unligated, indicating the transporter cannot be trapped under these conditions (**Figure 2A, bottom panels**). As prior work has shown EcMsbA is active with Mn^2+^,^30^ we conducted a series of functional assays using various divalent metals to identify conditions suitable for trapping PaMsbA (**Figure 2B**). Mn^2+^, Co^2+^, Cu^2+^, and Ni^2+^ support ATPase activity of EcMsbA and PaMsbA but to different extents and, in some cases, at comparable levels to that observed for Mg^2+^. Remarkably, Zn^2+^ significantly enhances the activity of PaMsbA by approximately four-fold compared to Mg^2+^. We also found ATPase activity was dependent on the concentration of Zn^2+^ (**Figure 2C**). At 1 mM of Zn^2+^ the activity is 52 ± 3 nmol Pi min^-1^ mg^-1^, which is within the range reported for EcMsbA.^36^ By substituting Mg^2+^ with Zn^2+^ in the vanadate trapping step, it becomes possible to effectively trap PaMsbA. The mass spectrum shows the majority of PaMsbA is now bound to adenosine 5’-diphosphates (ADPs), vanadates and Zn^2+^ ions, resulting in an experimental mass shift of 1145 ± 5 Da (**Figure 2D**). These results highlight the importance of Zn^2+^ in PaMsbA activity along with its importance in trapping the transporter with vanadate.

**Figure 2.**
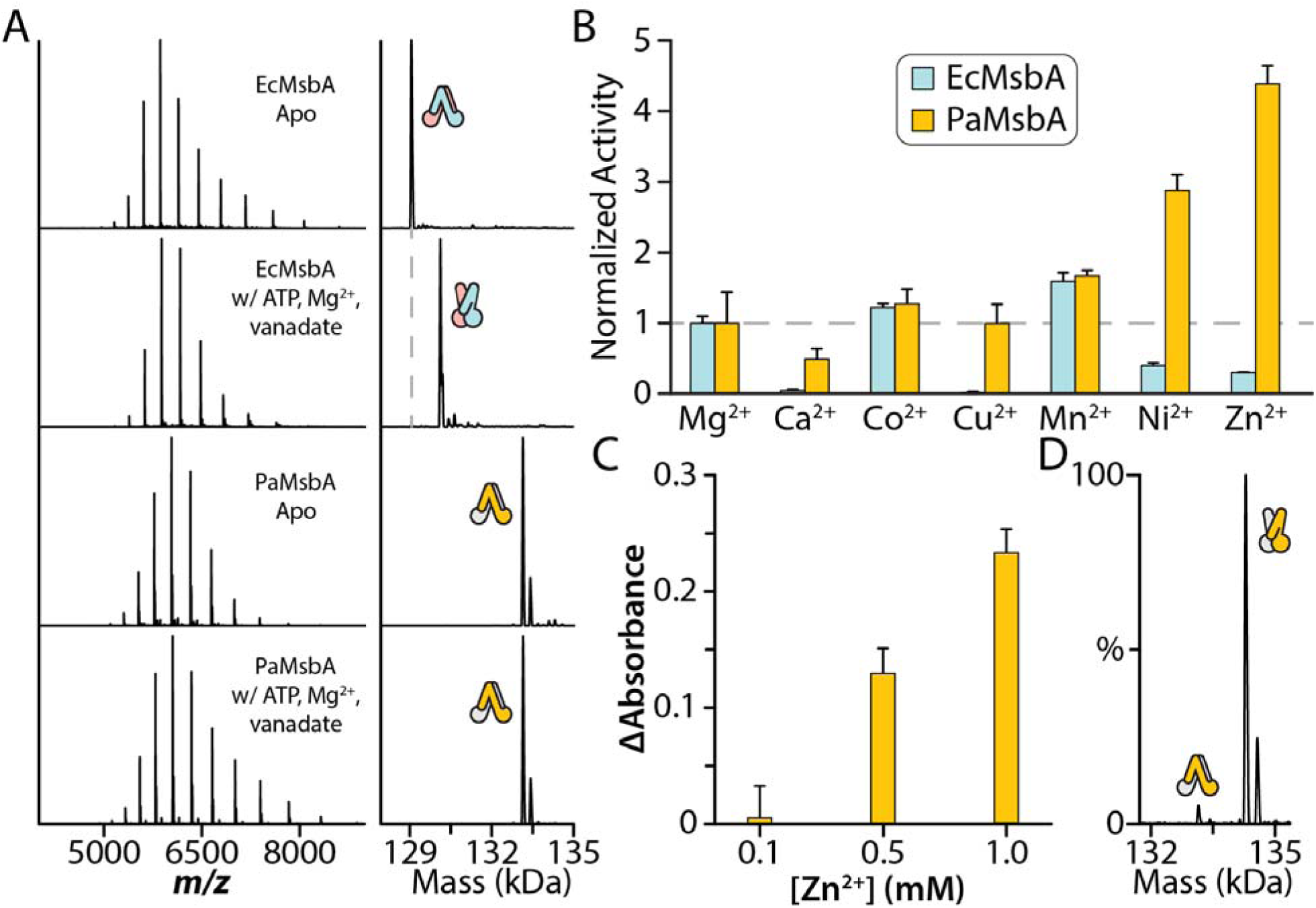
Selective stimulation of PaMsbA ATPase activity by divalent cations. A) Native mass spectra of EcMsbA and PaMsbA, and after incubating with ATP, Mg^2+^ and vanadate. EcMsbA is succesfully trapped with vanadate whereas PaMsbA is not trapped. B) ATPase activities of 0.5 µM EcMsbA or PaMsbA with 1 mM of different divalent cations and 1 mM of ATP. Activities are normalized to the corresponding activity with Mg^2+^. C) ATPase activity of PaMsbA in the presence of 1 mM ATP and different concentrations of Zn^2+^ incubated for ten minutes. Plotted is the change absorbance using a malachite assay. D) The native mass spectrum and the deconvolution of PaMsbA trapped by ADP, Zn^2+^ and vanadate. Reported are mean and standard deviation (*n* = 3).

To further validate our findings, we employed inductively coupled plasma mass spectrometry (ICP-MS) to analyze a subset of metals for EcMsbA and PaMsbA before and after trapping (**Table S2**). For EcMsbA, we observed an increase in Mg²⁺ concentration in the trapped sample. Consistent with our previous report, EcMsbA co-purified with Cu²⁺ but not Zn²⁺. Interestingly, PaMsbA co-purifies with Zn²⁺, and the concentration of this divalent cation was higher in the trapped state. In contrast, the Mg²⁺ levels remained consistent between PaMsbA samples, providing additional evidence that Zn^2+^ replaces Mg^2+^ in the active site for trapped PaMsbA.

### Determination of lipid binding affinity to vanadate trapped PaMsbA

To determine the selectivity of lipid binding to PaMsbA in an outward facing conformation, a series of lipid titrations were conducted using PaMsbA trapped with vanadate, ADP, and Zn^2+^ (**Figure 3**). Analogous experiments were performed to determine K_D_s for lipids binding to vanadate trapped PaMsbA (**Figure 3D and S4-S5**). In general, the lipid binding profiles for vanadate trapped PaMsbA are significantly different compared to the apo transporter. For example, the binding affinity of the first POPA molecule was enhanced (K_D1_=2.5 ± 0.3 μM) in comparison to untrapped PaMsbA (K_D1_=13.6 ± 0.9 μM) (**Figure 3B-D and Table S1 and S3**). In addition to POPA, the binding affinity of TOCDL significantly increased by six-fold, decreasing from 2.4 μM to 0.4 μM for K_D1_. However, unlike EcMsbA, the binding affinity of KDL did not significantly change when the transporter is trapped in an outward facing conformation (**Table S1 and S3**).

**Figure 3.**
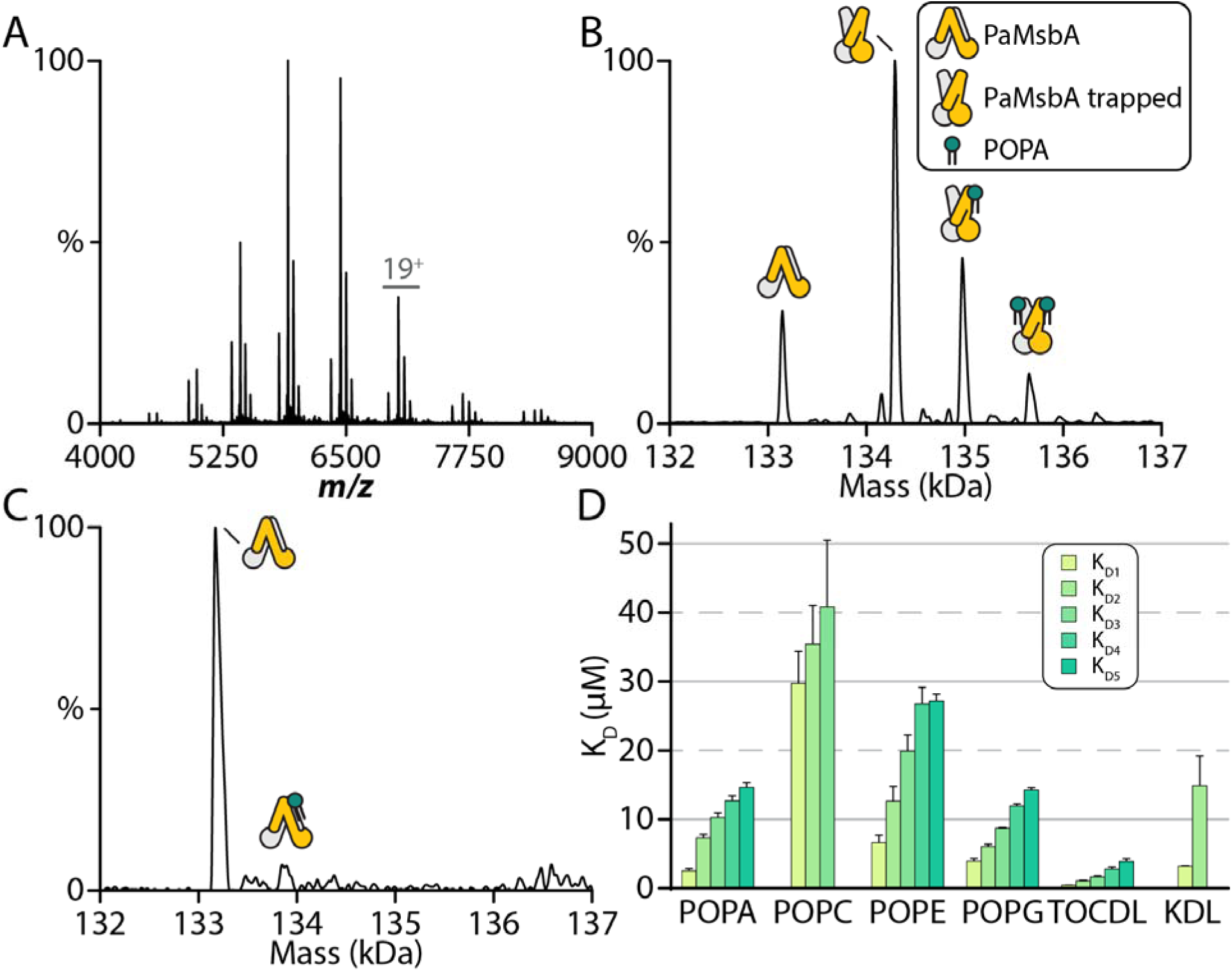
Determination of lipid binding affinities for PaMsbA trapped with vanadate. A) Native mass spectrum of 0.19 µM PaMsbA trapped with vanadate and in the presence of 1 µM POPA. B) Deconvolution of the mass spectrum shown in panel A. C) Deconvoluted mass spectrum of a mixture containing 0.3 µM PaMsbA and 1 µM POPA. The untrapped transporter binds less tightly to POPA. D) K_D_s for different lipids binding to vanadate trapped PaMsbA. Reported are mean and standard deviation (*n* = 3).

### Vanadate-trapped PaMsbA structures determined by cryoEM

To better understand the role of Zn^2+^ in activating PaMsbA, we set out to determine the structure of the transporter trapped in the presence of ATP, vanadate, and Zn^2+^. After screening several conditions for cryoEM studies, we found the trapped transporter solubilized in the dodecylmaltoside (DDM) detergent showed well distributed particles. From this dataset, we determined two cryoEM structures of the transporter in different conformations (**Figure S6 and Table S4-S5**). The first structure adopts an occluded, outward-facing conformation that was determined to a resolution of 2.98 Å (**Figure 4 and S7**). At this resolution, we were able to model the entire transporter with exception of a flexible loop (residues 68-73) connecting TM1 and TM2 and a disordered portion of the C-terminus (residues 592-603) (**Figure 4B**). Lipid-like density is observed that lies in the TMD near the N-terminus (**Figure 4C**). Four tube-like densities protrude into hydrophobic pockets and connected to a headgroup with a planar density. As the sample for cryoEM did not include a nonyl glucoside wash, which we found was necessary to produce well-resolved mass spectra (as shown in Figure 1), this density likely corresponds to a co-purified lipid, but detergent cannot be ruled out. The identity of molecule at this site is unknown but the planar density for the headgroup suggests it may correspond to an LPS precursor, such as lipid A. Moreover, well defined density enabled modeling of the ADP, vanadate, and Zn^2+^ bound at the NBDs (**Figure 4D**). Zn^2+^ is bound at a position consistent with that of Mg^2+^ in other transporters. Binding of Zn^2+^ is supported by native MS results showing the presence of Zn^2+^ is required to trap the transporter with vanadate (**Figure 2**). The distance between Zn^2+^ and S397 is ∼1.9 Å, consistent with a distance reported for this interaction.^39^ While the data is consistent with Zn^2+^ binding, there remains the possibility that Mg^2+^ or mixed metals can be bound in the NBDs. ADP and vanadate are coordinated by a series of conserved residues that is reminiscent to our recently reported structure of vanadate-trapped EcMsbA structure (**Figure 4D**).^34^ The occluded, outward-facing conformation is similar to the previously reported structure of EcMsbA (PDB 7BCW),^40^ which shares 40% sequence identity with PaMsbA, with a root mean square deviation of ∼2 Å.

**Figure 4.**
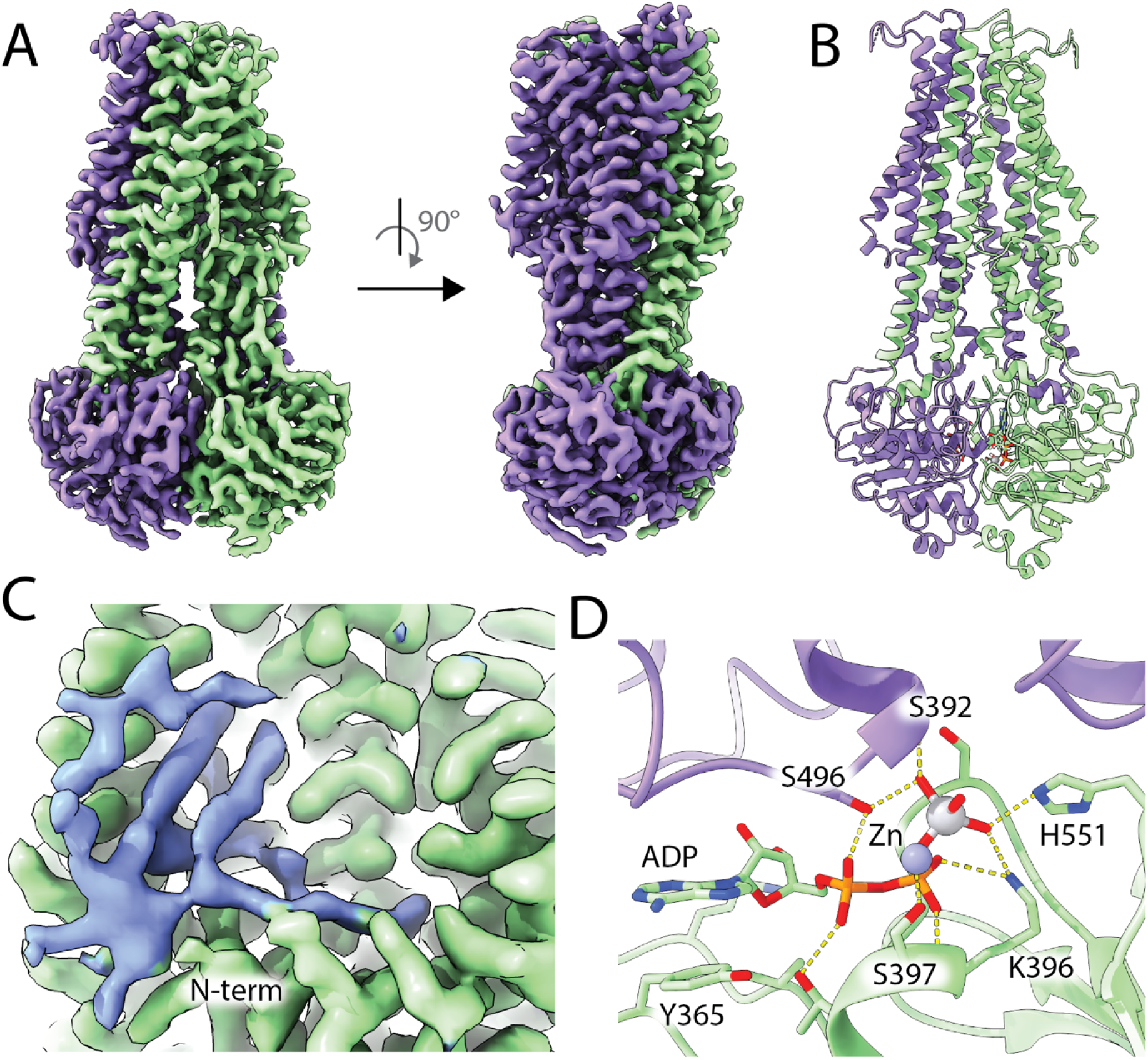
Structure of PaMsbA in an occluded, outward-facing conformation. A) CryoEM density map of PaMsbA as viewed from the perspective of the membrane plane. B) PaMsbA structure shown in cartoon representation. ADP, vanadate, and Zn^2+^ bound in the NBDs are shown in stick representation. C) View of lipid density (shown in blue) located near the N-terminus (N-term). Tube-like density penetrates a hydrophobic surface located in the TMD (shown in green). D) View of the NBD bound ADP, vanadate, and Zn^2+^. Interacting residues are shown in stick representation and bonds are shown as dashed yellow lines.

The other structure of PaMsbA was determined from the same dataset to a resolution of 2.72 Å (**Figure 5, S6 and S8**). In this structure, PaMsbA adopts an open, outward-facing conformation (**Figure 5B**). Like PaMsbA in the occluded state, the outward-facing structure shows well defined density for ADP, vanadate and Zn^2+^ bound in the NBDs. The coordination of these ligands is identical in both structures of PaMsbA. Aligning the open, outward-facing structure of PaMsbA with EcMsbA in a similar conformation (PDB 8DMM) reveals several differences (**Figure 5C**). More specifically, the PaMsbA structure adopts a more open conformation of the TMDs by about ∼6 Å (**Figure 5D**). Like the occluded PaMsbA structure, lipid-like density is observed in the TMD near the N-terminus (**Figure S9**). Additional density is observed near the bottom of the splayed TMDs, wedged between TM1 and TM3. DDM was modeled into this density and the detergent headgroup is coordinated by several residues (**Figure 5E and S8E**).

**Figure 5.**
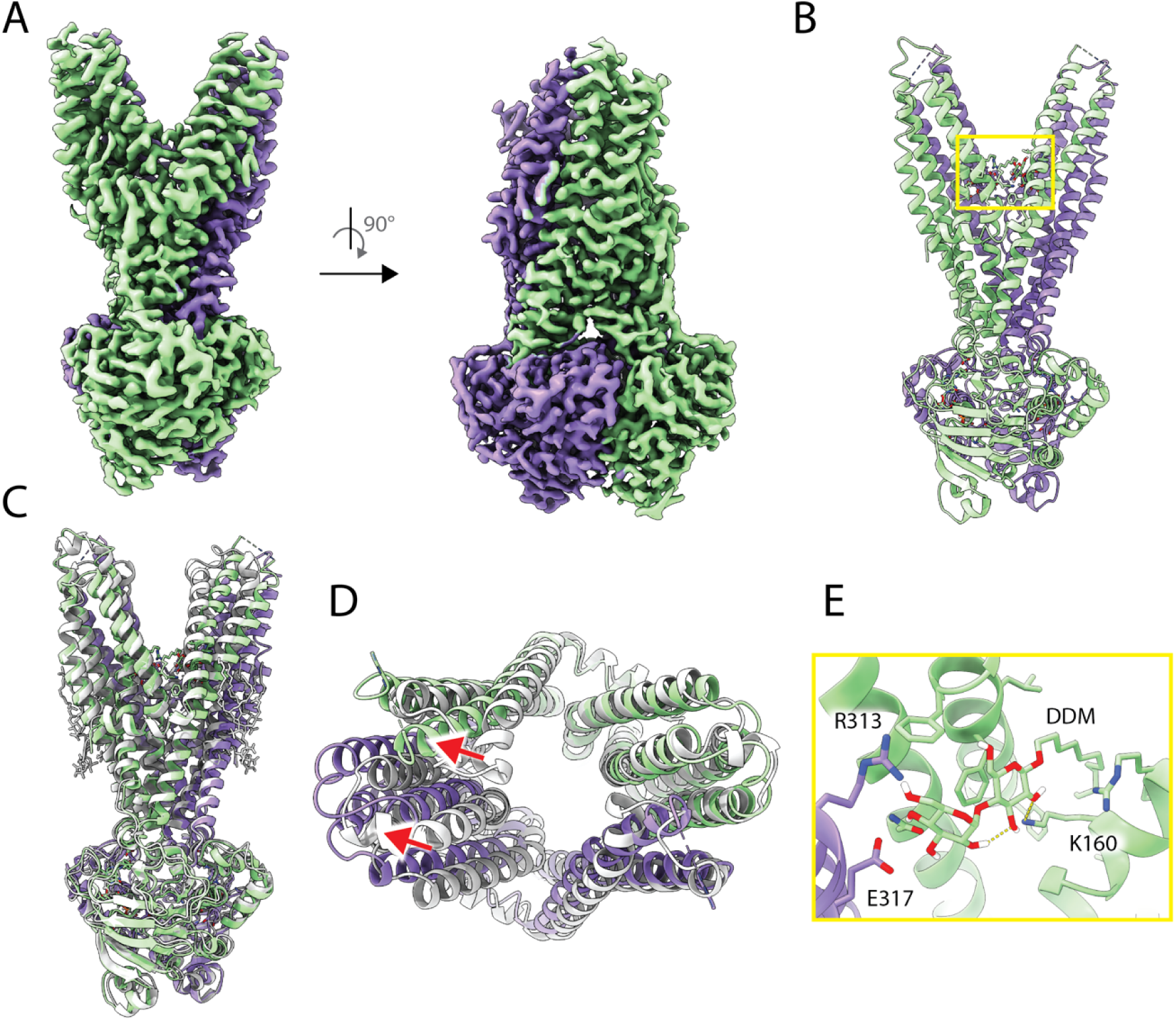
Open, outward-facing structure of PaMsbA. A) Two views of the cryoEM density map of PaMsbA from the perspective of the membrane plane. B) PaMsbA structure shown in cartoon representation. Bound ligands are shown in stick representation. C) Alignment of PaMsbA with a previous structure (PDB 8DMM) shown in silver. The structures were aligned to a single subunit, chain A. D) View from the periplasmic space of the aligned structures. PaMsbA is more open and highlighted by red arrows. E) View of DDM and interacting residues. Molecules are shown in stick representation with dashed yellow lines to indicate bonds.

To pinpoint possible Zn^2+^ binding sites, we examined the two PaMsbA structures. Interestingly, we found a triad of histidine residues formed by H135, H139, and H221 (**Figure 6A**). The density maps for both structures suggest that the histidine residues may be coordinating Zn^2+^ (**Figure 6A**). Moreover, the distance between H135 and H139 (Cγ to Cγ) range from 5 to 6 Å, an anticipated distance for these sidechains bridged through coordination to Zn^2+^.^41^ The triad of histidine residues in PaMsbA differ in EcMsbA with corresponding positions occupied by glutamine, serine, and threonine, specifically at positions 135, 139, and 221, respectively.

**Figure 6.**
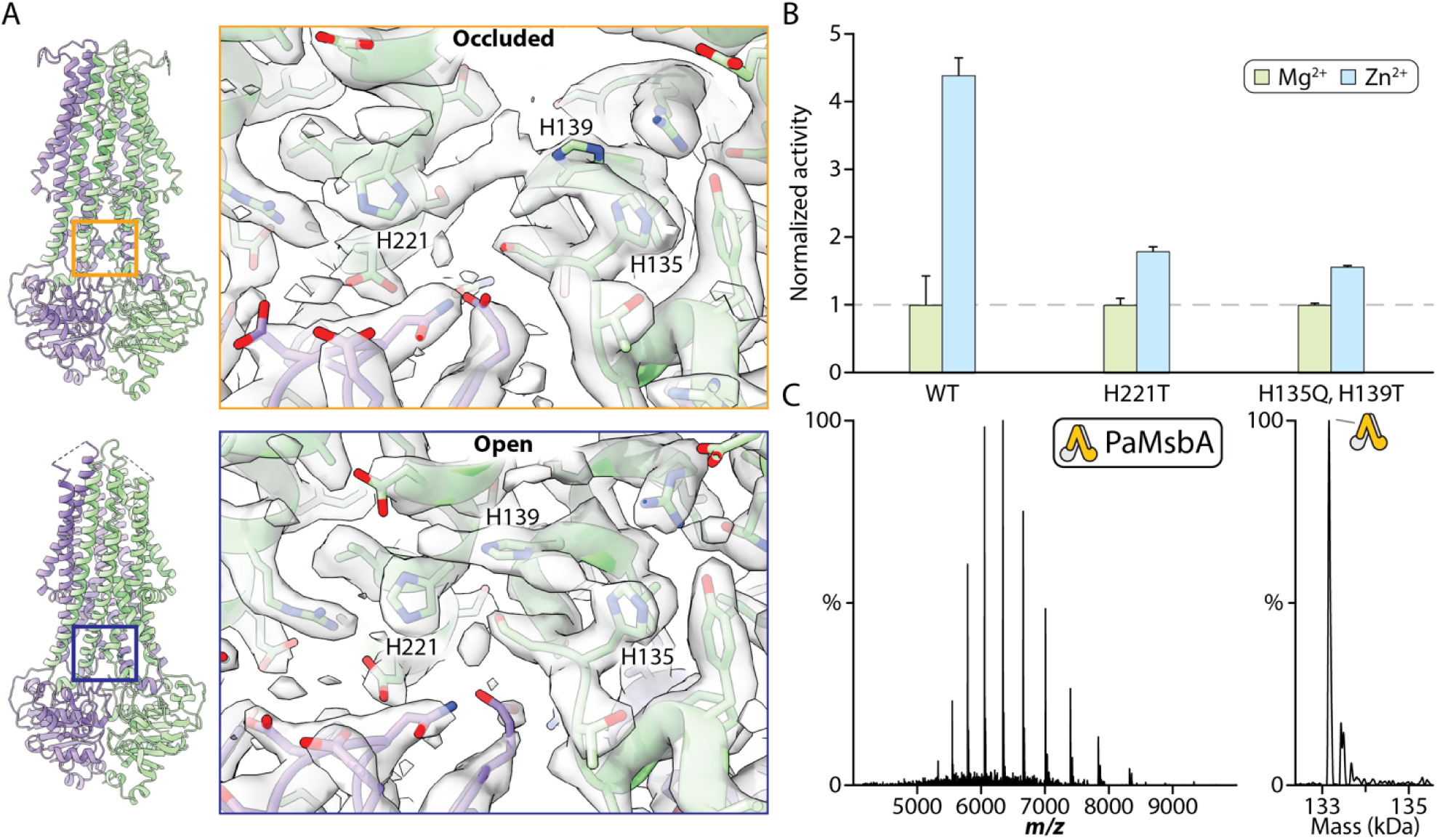
Identification and functional implications of a histidine triad in PaMsbA. A) Structure of PaMsbA in cartoon with box highlighting position of histidine residues. Shown to the right are views of the histidine residues in the two conformations of PaMsbA. Residues are shown in stick representation and cryoEM density map is shown as a transparent surface. B) ATPase activities of 0.5 µM PaMsbA and mutants with 1 mM of ATP and 1 mM of Mg^2+^ or Zn^2+^. Activities are normalized to the corresponding activity with Mg^2+^. Reported are the mean and standard deviation (*n* = 3). C) Native mass spectrum of PaMsbA under conditions to trap with vanadate and Zn^2+^ but in the presence of DTPA. Chelation of Zn^2+^ prevents vanadate-trapping of PaMsbA.

### Probing the Zn^2+^ binding site in PaMsbA

Several mutations were incorporated into PaMsbA to assess the consequences of histidine residues – H135, H139, and H221 – in binding Zn^2+^. As demonstrated above, monitoring the ATPase activity of PaMsbA showed Zn^2+^ significantly enhanced the activity of the wildtype transporter compared to Mg^2+^ (**Figure 2B and 6B**). To match the sequence of EcMsbA, we introduced the single mutation of H221T and the double mutation of H135Q and H139T. The mutant transporters show significant impairment of the Zn^2+^ stimulation of ATPase activity (**Figure 6B**). We conclude that the triad of histidine residues in PaMsbA are important for stimulation of the transporter by Zn^2+^. In addition, we treated PaMsbA under trapping conditions (**Figure 2A**) but in the presence of diethylenetriamine pentaacetate (DTPA), a zinc chelator (**Figure 6C**). The results reveal trapping of PaMsbA is impaired, providing additional evidence that Zn^2+^ is required for ATPase activity and trapping with vanadate.

## Discussion

MsbA and other transporters are known to require a metal cofactor, typically Mg^2+^, to hydrolyze ATP and carry out function.^32,42,43^ Here, we uncover an unexpected role of Zn^2+^ in function along with its requirement to trap the transition state of PaMsbA with vanadate. A previous report has shown EcMsbA is more active in the presence of Mn^2+^ than Mg^2+^,^30^ which we also observe a modest enhancement in activity. However, the ATPase activity of PaMsbA is four-fold more active in the presence of Zn^2+^ compared to Mg^2+^. While not much has been reported regarding Zn^2+^ stimulating MsbA activity, one report found Zn^2+^ stimulated the activity of multidrug resistance-related protein 2 (Mrp2), p-glycoprotein (P-gp), and breast cancer resistance protein (Bcrp).^44^ Earlier structural studies reported the inability to trap BtuCD with a non-hydrolysable ATP analogue in the presence of Mg^2+^.^45^ Based on this work, it is possible BtuCD could not be trapped due to the choice of the metal co-factor. In short, these results illustrate the powerful utility of native MS to identify (and optimize) conditions to trap PaMsbA for structural and biochemical studies.

Biophysical characterization of PaMsbA-lipid interactions by native MS begins to define the molecular requirements that support lipid binding to PaMsbA in different conformational states. Like our previous study on EcMsbA,^34^ we find that some but not all lipids display an altered affinity for the transporter in an inward-facing versus outward-facing conformations. In particular, the binding of POPA, POPG, and TOCDL is enhanced by more than two-fold, in some cases as high as six-fold. These results suggest the different conformations of the transporters, which results in restructuring of the TMD, has unique lipid binding pockets. While we focus here on lipids with similar acyl chains, our recent work on TRAAK, a two-pore domain potassium channel, has shown a remarkable selectivity for the lipid acyl chains but also the fatty acid linkage.^46^ It remains unclear how the chemistry of lipid tails will influence binding affinities and warrants further study.

The structures of PaMsbA reveal not only two outward-facing conformations (open and occluded)of the transporter when trapped in the transition state by vanadate but also a triad of His residues. Open and occluded outward-facing conformations have been previously reported for EcMsbA^24,29,34,40^ that are largely similar to the PaMsbA structures. A notable exception is the TMDs are more open in the PaMsbA structure. Nearly equal number of particles were selected from the dataset for both conformations, suggesting these two states are in equilibrium. In addition, lipid-like density is observed near the N-terminus with tube-like density interacting within a hydrophobic pocket of the TMD in both conformations. However, the identity and role of this lipid in the PaMsbA transport cycle is unclear.

We show PaMsbA activity is stimulated by Zn^2+^ and the two structures of PaMsbA reveal a triad of histidine residues (**Figure 6 and S10**). Functional studies implicate the histidine residues are associated with the Zn^2+^ induced stimulation of ATPase activity. One of these histidine residues is situated within a loop region positioned between one of the coupling helices and TM3, and another found on TM3. The third histidine is located on TM4. While TM3 and TM4 associate in outward-facing conformations of MsbA, in open, inward-facing structures these helices are separated with distances depending on the openness of the transporter. For example, the open, inward-facing EcMsbA structure (PDB 8DMO) are separated ∼53 Å (D117 Cα to Cα). The ATPase activity of EcMsbA is known to be stimulated by LPS, which is thought to promote dimerization of the NBDs.^36–38^ It is intriguing to consider whether Zn^2+^ plays a similar role by facilitating the transition of MsbA from an open to a closed structure, which the driving force is mediated by specific coordination of Zn^2+^.

Taken together, our combined structural and functional data offer insights into the lipid binding preferences of PaMsbA and reveal essential residues linked to the enhancement of ATPase activity by Zn^2+^. We further elucidate structural differences between PaMsbA and EcMsbA, revealing an open, outward-facing conformation that is more open. The results presented here suggest a mechanism by which Zn^2+^ promotes dimerization of the NBDs of PaMsbA through coordination with specific histidine residues located within the TMD, leading to enhanced ATPase activity.

## Materials and Methods

### MsbA expression constructs

Expression constructs for PaMsbA and mutants were essentially performed as previously described for EcMsbA.^34,47^ In detail, the PaMsbA gene (UniProt Q9HUG8) was codon optimized for *E. coli*, synthesized (Twist Bioscience), and cloned into a modified pCDF-1b plasmid (Novagen) to produce PaMsbA with an N-terminal His6 fusion protein that includes a TEV protease cleavage site. Point mutations were introduced using the Q5 Site-Directed Mutagenesis Kit (New England Biolabs) according to manufacturer’s instructions. All expression constructs were verified through DNA sequencing.

### Protein expression and purification

PaMsbA and mutants were essentially expressed and purified as previously described for EcMsbA.^34,47^ In detail, *E. coli* (DE3) BL21-AI competent cells (Invitrogen) were transformed with PaMsbA expression plasmids. A single colony was selected to inoculate 50 mL of LB medium and cultured overnight at 37 °C with shaking at 200 rpm. Next day, terrific broth (TB) media was inoculated with the overnight culture and incubated at 37 °C until the OD_600_ reached 0.6-1.0. After that, the culture was supplemented with a final concentration of 0.5 mM IPTG (isopropyl β-D-1-thiogalactopryanoside) and 0.2% (w/v) arabinose to induce protein expression. After overnight expression at 25 °C and shaking at 200 rpm, the TB cultures were harvested by centrifugation at 4000 x g for 10 minutes and the resulting pellet was resuspended in lysis buffer (20 mM TRIS, 300 mM NaCl and pH at 7.4 at room temperature). The resuspended cells were centrifuged to wash the cells. The pellet was resuspended in lysis buffer before being lysed by four passages through a Microfluidics M-110P microfluidizer operating at 25,000 psi with reaction chamber emersed in an ice bath. The lysate was centrifuged at 20,000 x g for 25 minutes and the resulting supernatant was centrifuged at 100,000 x g for 2 hours to pellet membranes. Resuspension buffer (20 mM Tris, 150 mM NaCl, 20% (v/v) glycerol, pH 7.4) was used to homogenize the resulting pellet and 1% (w/v) DDM was added to extract membrane proteins with incubation overnight at 4 °C. The extraction was centrifuged at 20,000 x g for 25 minutes to pellet insoluble material. The resulting supernatant was supplemented with 10 mM imidazole and filtered with a 0.45 µm syringe filter prior to purification by immobilized metal affinity chromatography. The extraction containing solubilized PaMsbA was loaded onto a column packed with 2.5 mL Ni-NTA resin pre-equilibrated in NHA-DDM buffer (20 mM TRIS, 150 mM NaCl, 10 mM imidazole, 10% (v/v) glycerol, pH 7.4 and supplemented with 2 x the critical micelle concentration (CMC) of DDM). The column was then washed with 5 column volumes (CV) of NHA-DDM buffer, 10 CV of NHA-DDM buffer supplemented with 2% (w/v) nonyl-ß-glucoside (NG) for delipidation followed by 5 CV of NHA-DDM buffer. The immobilized protein was eluted by the addition of 2 CV of NHB-DDM buffer (20 mM TRIS, 150 mM NaCl, 250 mM imidazole, 10% (v/v) glycerol, 2 x CMC of DDM, pH 7.4). The eluted protein was pooled and desalted using HiPrep 26/10 desalting column (GE Healthcare) pre-equilibrated in desalting buffer (NHA-DDM with imidazole omitted). TEV protease (expressed and purified in-house) and 10 mM 2-mercaptoethanol (BME) was added to the desalted sample and incubated overnight at room temperature. The sample was passed over a pre-equilibrated Ni-NTA column and the flow-through containing the cleaved protein was collected. The pooled protein was concentrated using a centrifugal concentrator (Millipore, 100 kDa) prior to injection onto a Superdex 200 Increase 10/300 GL (GE Healthcare) column equilibrated in 20 mM TRIS, 150 mM NaCl, 10% (v/v) glycerol and 2 x CMC C_10_E_5_. Peak fractions containing dimeric PaMsbA were pooled, flash frozen in liquid nitrogen, and stored at −80 °C prior to use.

### Preparation of PaMsbA for native MS studies

PaMsbA samples were buffer exchanged into 200 mM ammonium acetate supplemented with 2 x CMC of C_10_E_5_ using a centrifugal buffer exchange device (Bio-Spin, Bio-Rad) for native MS studies. To prepare vanadate-trapped PaMsbA, 10 mM ATP and 10 mM ZnCl_2_ were added to PaMsbA sample and incubated at room temperature for 10 minutes. Ammonium vanadate (pH adjusted to 10) was added to reach final concentration of 1 mM and incubated at 37 °C for 10 minutes. The sample was then buffer exchanged as described above.

### Native mass spectrometry

Samples were loaded into gold-coated glass capillaries made in-house^48^ and introduced into a Thermo Scientific Exactive Plus Orbitrap with Extended Mass Range (EMR). For native mass analysis, the instrument was tuned as follows: source DC offset of 10 V, injection flatapole DC to 8.0 V, inter flatapole lens to 4, bent flatapole DC to 3, transfer multipole DC to 3 and C trap entrance lens to 0, trapping gas pressure to 6.0 with the in-source CID to 65.0 eV and CE to 100, spray voltage to 1.70 kV, capillary temperature to 200 °C, maximum inject time to 200 ms. Mass spectra were acquired with a setting of 17,500 resolution, microscans set to 1 and averaging set to 100.

### Metal analysis of MsbA samples

Samples were analyzed for elemental composition at the Elemental Analysis Laboratory at Texas A&M University using inductively coupled plasma mass spectrometry (ICP-MS). A PerkinElmer NexION ICP mass spectrometer was employed, with operating parameters following previously established protocols.^34^

### Preparation of PaMsbA for cryoEM studies

The expression and purification of PaMsbA for cyroEM studies was slightly modified. The immobilized protein was not washed with buffer containing NG, and before injecting onto the Superdex 200 Increase 10/300 GL column equilibrated in GF buffer (20 mM TRIS, 150 mM NaCl and 2 x CMC DDM) PaMsbA was trapped by vanadate using the method described for native MS studies. Peak fractions containing dimeric PaMsbA were pooled and concentrated to 17 mg/ml measured by UV 280 nm. ATP, zinc chloride, and ammonium vanadate were added to PaMsbA sample at a final concentration of 100 μM. The resulting mixture was filtered through a 0.2-micron filter. 3 μl of the sample was applied to glow-discharged Quantifoil R1.2/1.3 Orthogonal Array with 300M Cu grids. The grids were then blotted at 4°C in a 100% humidity chamber and plunge-frozen in liquid ethane using the Vitrobot Mark IV (Thermo Fischer Scientific). Data collection was performed at the LBSD (Texas A&M cryoEM center) using a Titan Krios microscope operated at 300 keV equipped with a BioQuantum energy filter and a post-GIF K3 direct electron detector. The magnification was set to 105,000, corresponding to a pixel size of 0.832 Å. The dose rate was 14.634 e− per pixel per second, and a total dose of 50.73 e− per Å2 was fractionated over 40 frames. Data collection was conducted using the EPU (version 2.7) software with a defocus range of −1 μm to −2.4 μm.

### Model building, refinement, and validation

The AlphaFold model of PaMsbA^49^ was docked into the cryo-EM map using Chimera.^50^ The model was manually refined and ligands were modeled into the density using Coot.^51^ The final model was subjected to one round of real-space refinement using Phenix.^52^ The final model refinement, statistics, and geometry can be found in **Table S4 and S5**. Figures were generated using ChimeraX ^53^ and Pymol (Schrödinger LLC., version 2.1).

### PaMsbA activity assay

The ATPase activity of PaMsbA was determined using a malachite green assay.^34,54^ 500⍰nM PaMsbA, 1⍰mM metal chloride, and 1 mM ATP were used for the assay. Reactions were carried out at 37⍰°C for 10⍰min and quenched by the addition of malachite green solution containing a 3:1 mixture of 0.045% (w/v) malachite green and 4.2% (w/v) ammonium molybdate prepared in 4⍰N HCl, 0.04% (v/v) Triton X-100 (final concentration). Then 3.1% (w/v) sodium citrate was added (final concentration) to enhance the coloring reaction. The quenched reactions were incubated at room temperature for 30⍰min and absorbance at 650⍰nm was measured with a CLARIOstar plate reader (BMG LabTech). The ATP hydrolysis rate was calculated from the slope fitted to the absorbance collected at 0 min (blank) and 10 min reaction times.

## Author contributions

J.L., H.B., T.Z., and A.L. designed the research. J.L. and T.Z. expressed and purified proteins and performed experiments. J.L., H.B., G.Y., and A.L., performed structural biology experiments. J.L., H.B., T.Z., and A.L. drafted the manuscript with input from the other authors.

## Acknowledgements

We thank Bryan Tomlin in the Elemental Analysis Laboratory for elemental analysis. This work was supported by Welch Foundation (A-2106-20220331) and National Institutes of Health (NIH) (R01GM138863, R01GM139876 and RM1GM145416) awarded to A.L., and NIH (RM1GM149374) awarded to D.R.

## Competing Interests

The authors declare no competing financial interests.

## Data and materials availability

The PaMsbA structures have been deposited with accession codes EMD-44444 (PDB 9BD6) and EMD-44445 (PDB 9BD7). All other study data are included in the article and/or *SI Appendix*.

## Supplementary Materials

Figs S1 to S10

Tables S1 to S5

## Notes

### Competing Interest Statement

The authors have declared no competing interest.

