## Supplementary Information for "Molecular basis for the activation of *Pseudomonas aeruginosa* MsbA by Zn^2+^"

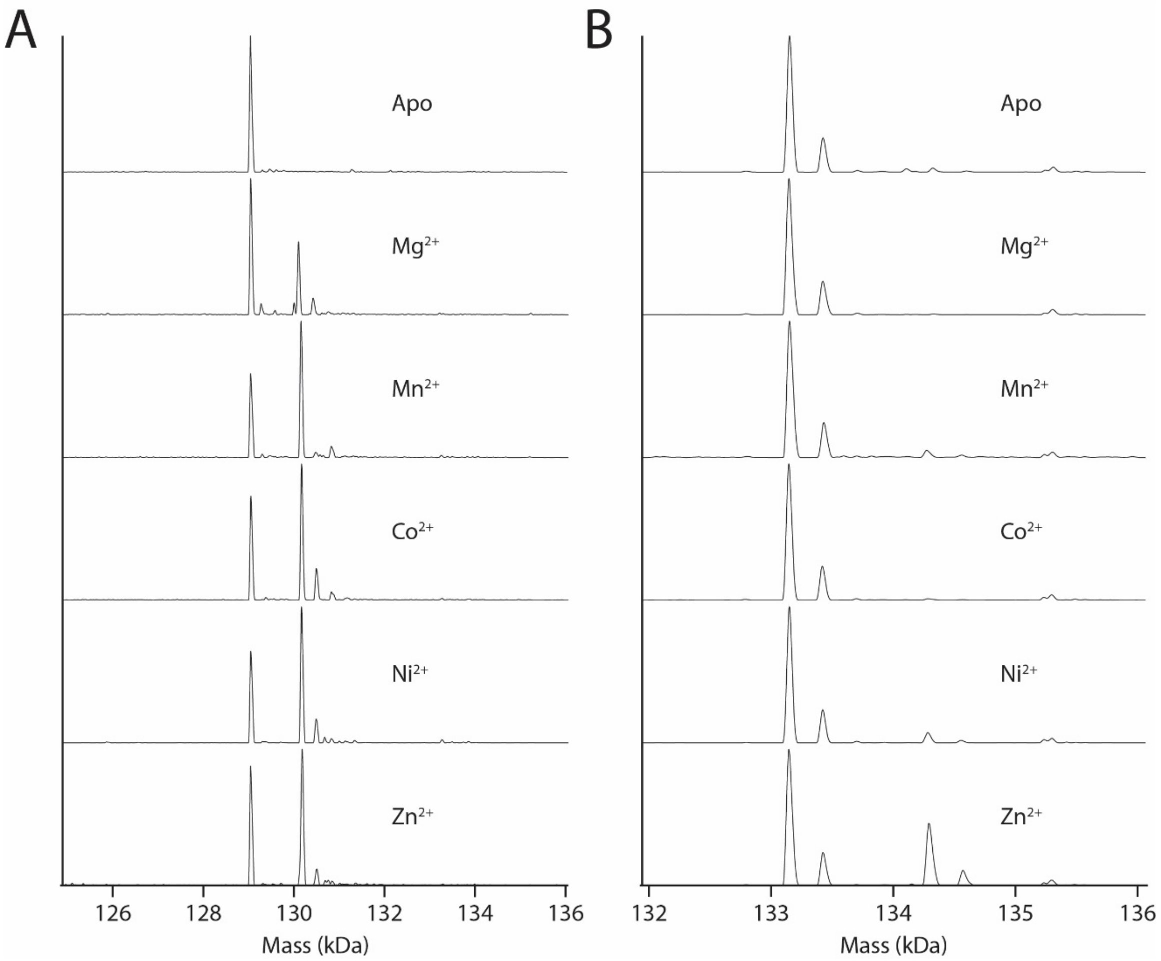


**Figure S1. Exploration of conditions to trap MsbA with vanadate.** Shown are deconvoluted native mass spectra of A) EcMsbA and B) PaMsbA containing a mixture of the transporter alone (added before native MS analysis) and after incubating in the presence of ATP and vanadate with different metals ions. Metal ions are denoted in the inset. All the divalent metals are capable of trapping EcMsbA with ADP and vanadate. However, $Zn^{2+}$ is the only metal ion capable of trapping PaMsbA with vanadate.

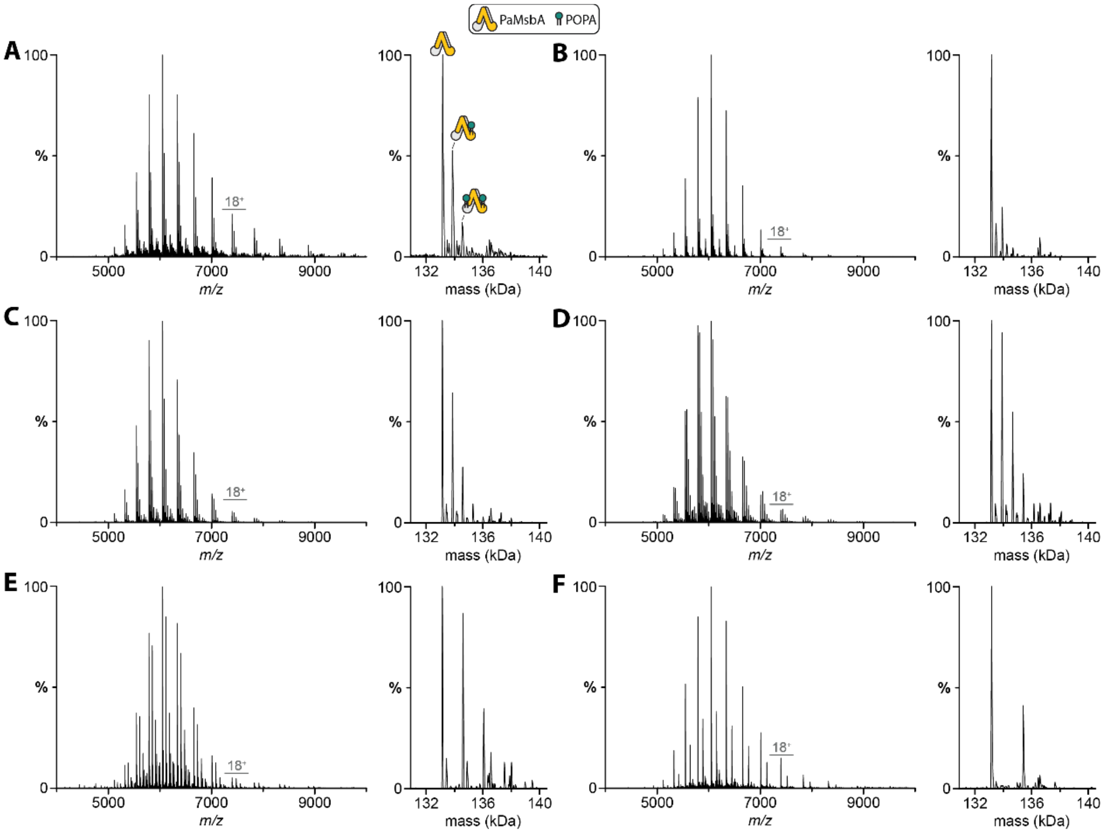


**Figure S2. Representative native and deconvoluted mass spectra for lipid binding to PaMsbA.** A) Mass spectra of PaMsbA (0.30 µM) mixed with 6 µM of POPA. B-E) PaMsbA (0.44 µM) mixed with B) 10 µM POPC, C) 10 µM POPE, D) 6 µM POPG, and E) 2 µM TOCDL. F) PaMsbA (0.25 µM) mixed with 1 µM of KDL.

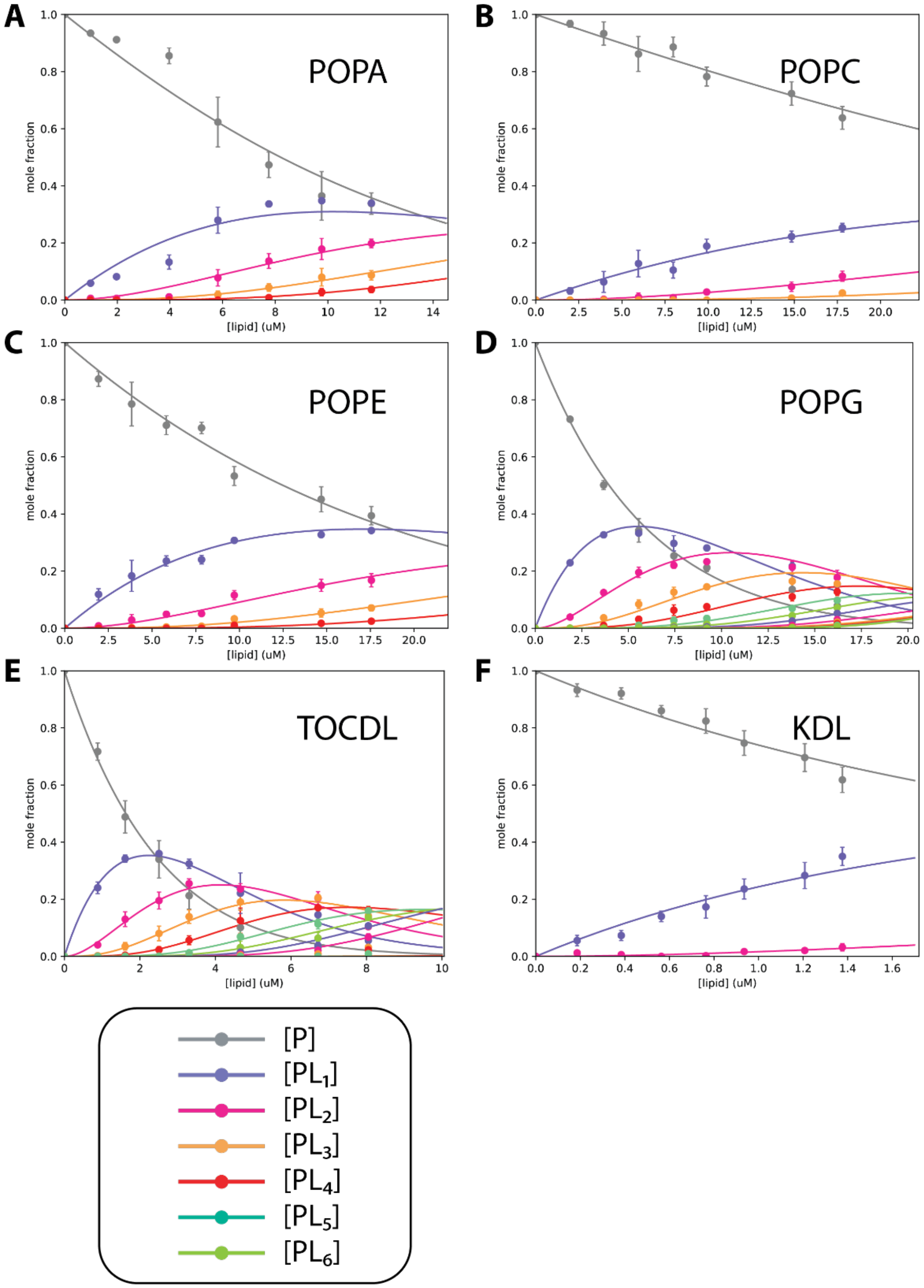


**Figure S3. Determination of equilibrium dissociation constants ($K_D$) for lipid binding to PaMsbA.** Plots of mole fraction for apo PaMsbA and bound to different number of lipids (dots) and resulting fit from a sequential lipid-binding model (solid lines). The different lipids are labelled. Reported are the mean and standard deviation (*n* = 3, biologically independent samples).

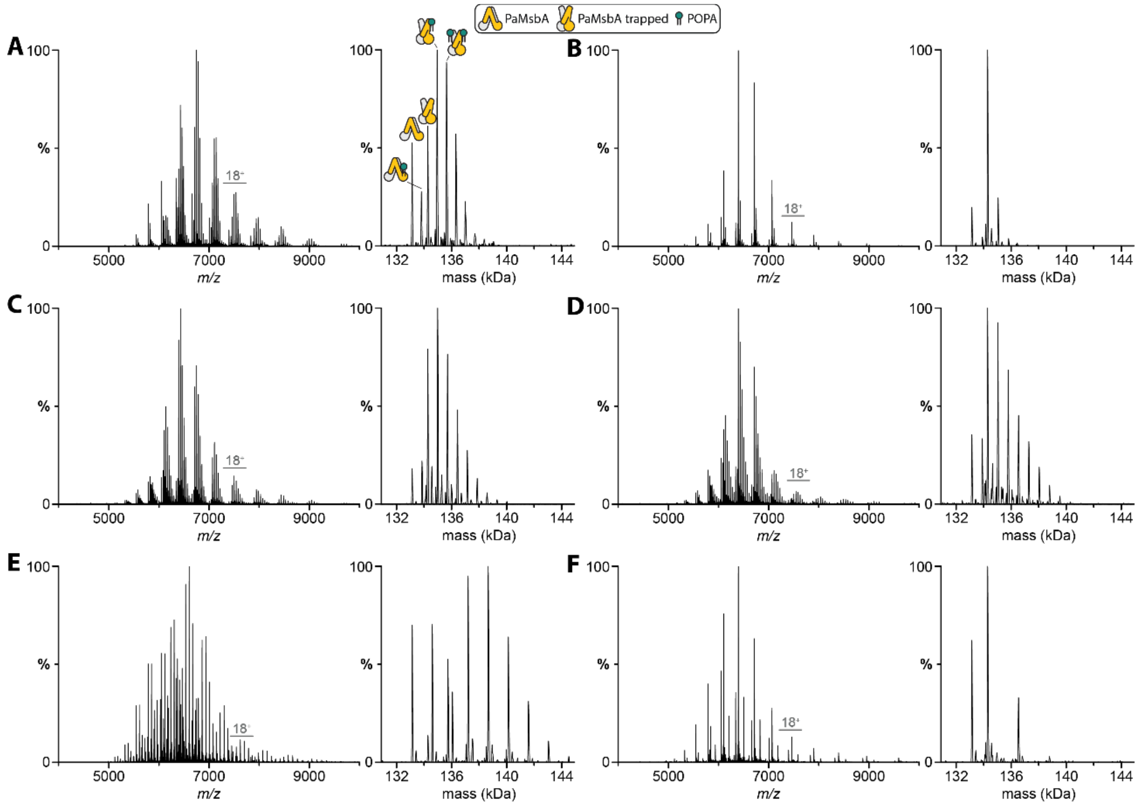


**Figure S4. Representative native and deconvoluted mass spectra for lipid binding to PaMsbA trapped by vanadate and Zn.** A) 0.19 µM of PaMsbA with 6 µM of POPA. B) 0.34 µM of PaMsbA with 10 µM of POPC, C) 0.23 µM of PaMsbA with 10 µM of POPE, D) 0.23 µM of PaMsbA with 5 µM of POPG, E) 0.22 µM of PaMsbA with 3 µM TOCDL. F) 0.24 µM of PaMsbA with 1 µM of KDL.

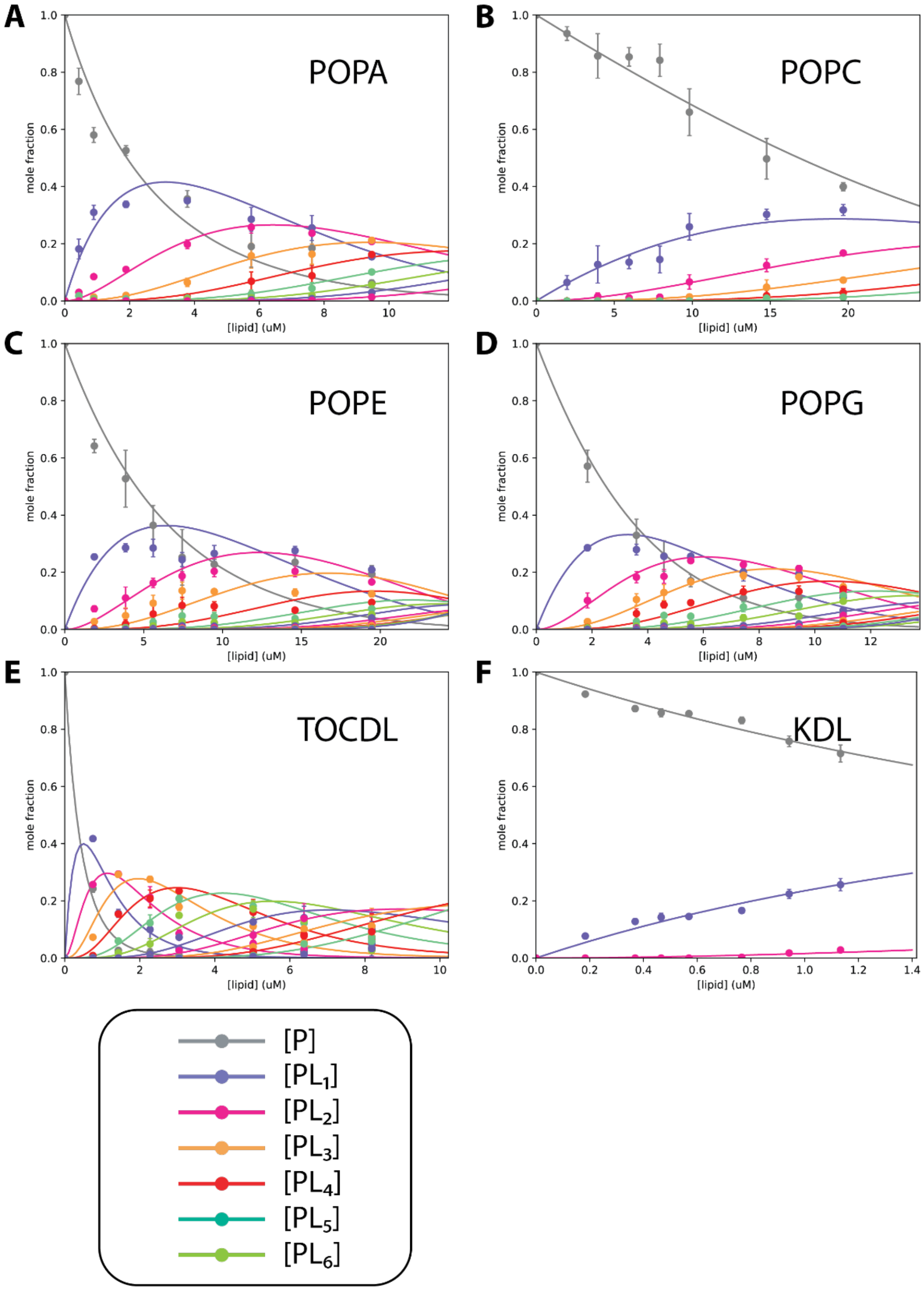


**Figure S5. Determination of $K_D$s for lipids binding PaMsbA trapped by vanadate and Zn.** Plots of mole fraction for apo PaMsbA bound to different number of lipids (dots) and resulting fit from a sequential lipid-binding model (solid lines). The different lipids are labelled. Reported are the mean and standard deviation (*n* = 3, biologically independent samples).

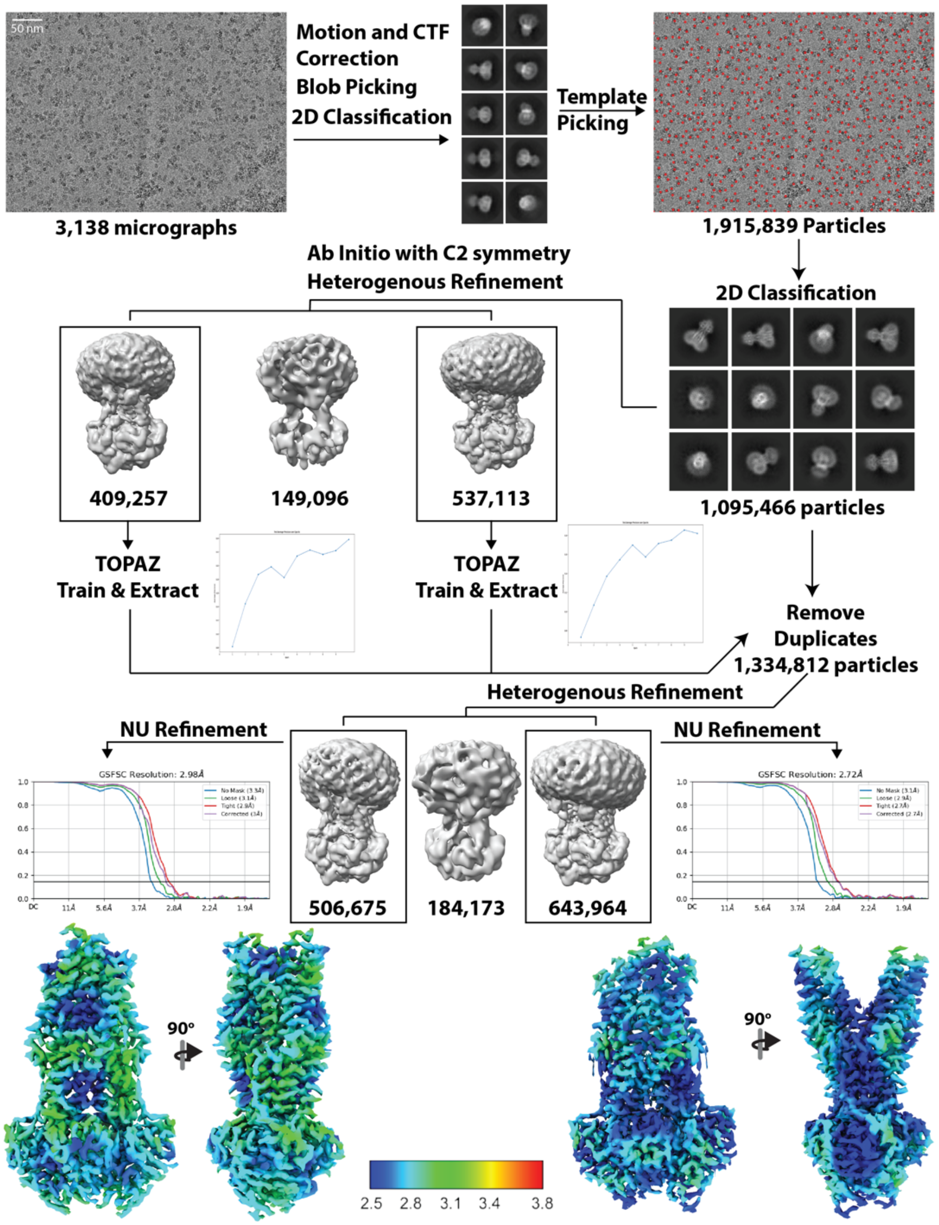


**Figure S6. CryoEM analysis of PaMsbA bound to ADP, vanadate, and $Zn^{2+}$.** Shown is a summary of the data processing workflow in cryoSPARC.[1] A motion-corrected micrograph is presented. Particle selection was performed by selection of 2D templates generated through 2D classification of picks using the blob picker. The figures shown depict representative 2D class averages with a box edge of 225 Å. Three models were generated through ab initio construction followed by heterogenous refinement. C2 symmetry was applied in this and subsequent steps. A subset of particles were used to train TOPAZ[2] extract particles from all micrographs. These were combined using the remove duplicate tool. This process resulted in more particles and were subjected to heterogenous refinement. Non-uniform refinement was performed on the two best models. Fourier shell correlation curves of the final reconstruction and model versus map are shown for the two resulting structures. The resolution of the reconstruction was determined by the FSC=0.143 criterion.

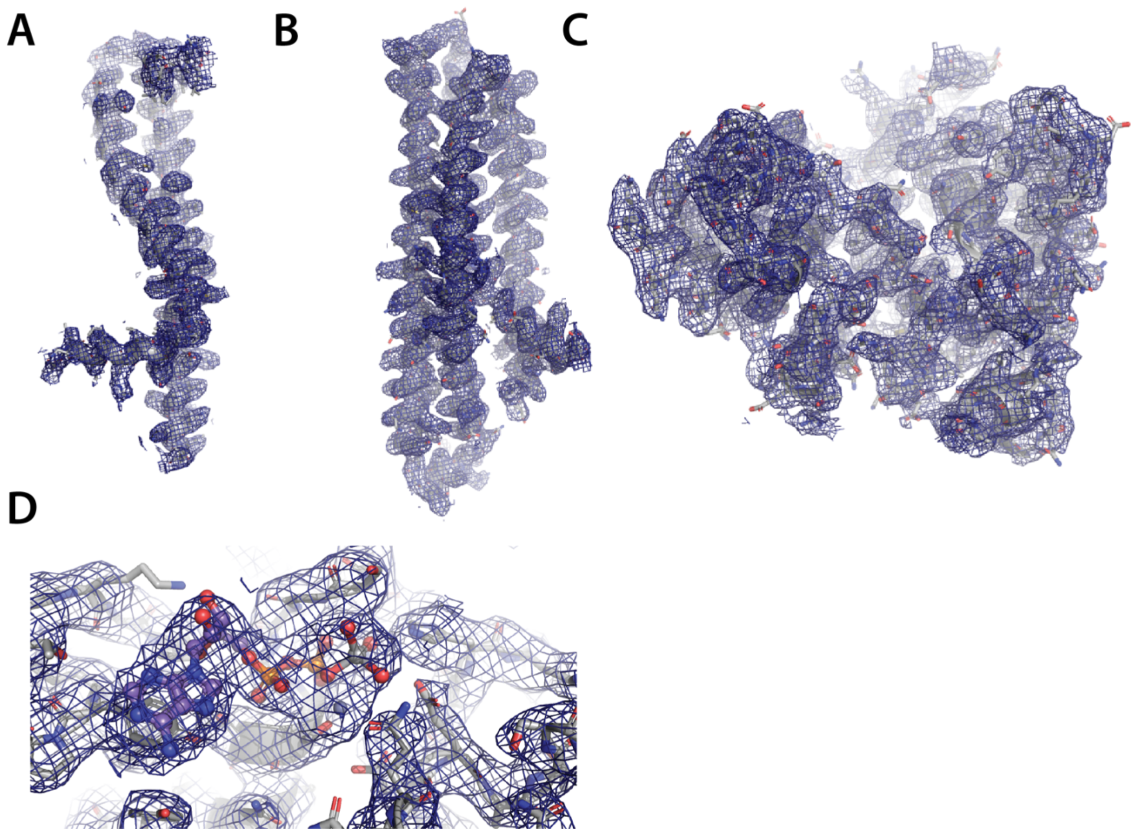


**Figure S7. CryoEM density of the occluded, outward-facing PaMsbA structure.** Shown are the density and atomic model for different regions of PaMsbA: (A) TM1-2, (B) TM3-6, (C) NBD, and (D) ADP-vanadate and $Zn^{2+}$. Figure was prepared using Pymol[3] and maps contoured at 5 sigma.

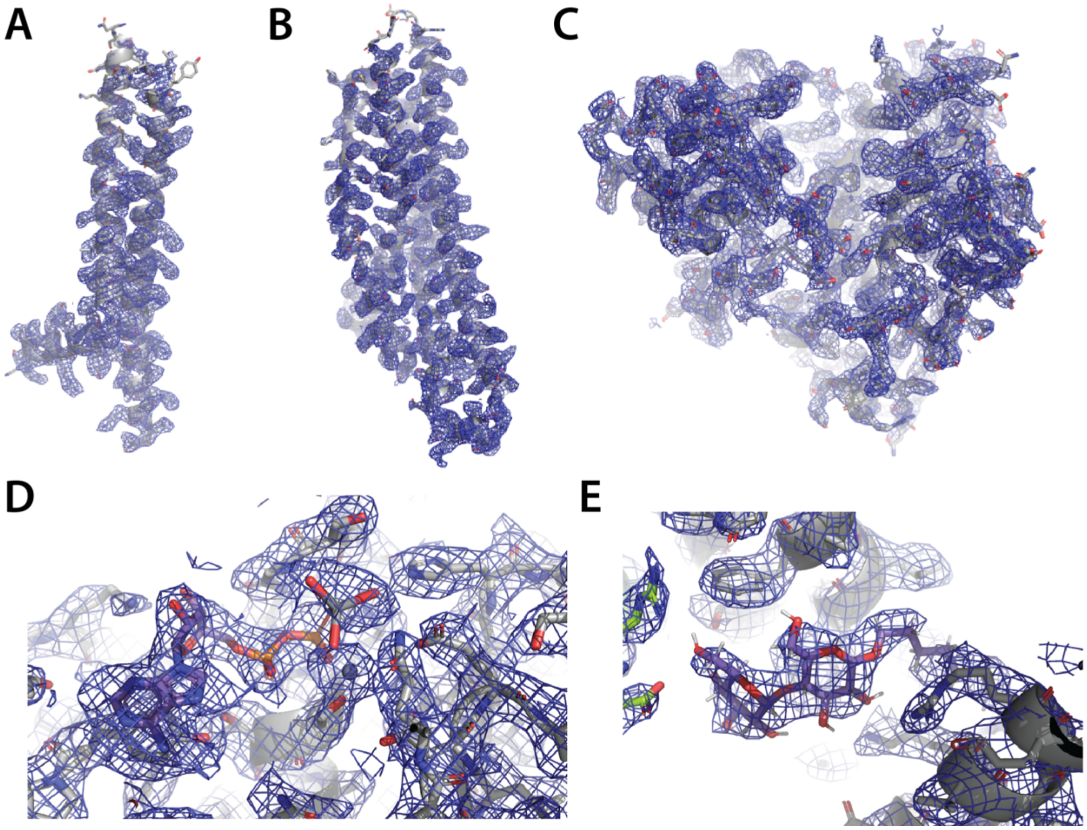


**Figure S8. CryoEM density of the open, outward-facing PaMsbA structure.** Shown are the density and atomic model for different regions of PaMsbA: (A) TM1-2, (B) TM3-6, (C) NBD, (D) ADP-vanadate and $Zn^{2+}$, and (E) DDM. Figure was prepared using Pymol[3] and maps contoured at 5 sigma.

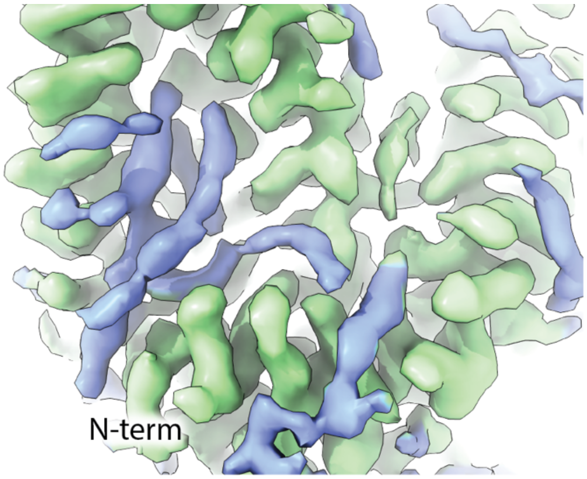


**Figure S9. Lipid-like density in the open, outward-facing PaMsbA structure.** Density for protein and lipid is shown in green and blue, respectively. Density for lipid acyl chains interact with the hydrophobic TMD. Shown in a similar orientation as Figure 4C.

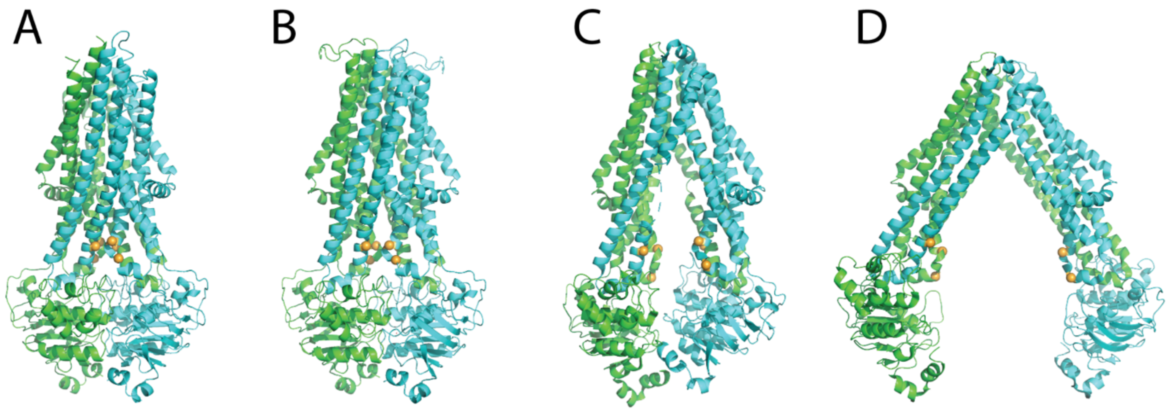


**Figure S10. Location of histidine residues in different conformations of MsbA.** Shown are the structures in cartoon representation of (a) open outward-facing (PaMsbA reported here), (b) occluded outward-facing (PaMsbA, reported here), (C) G907 bound closed inward-facing (EcMsbA, PDB 6BPL), and (d) open inward-facing (EcMsbA, PDB 8TSQ). The Calpha of His residues (135, 139, and 221) in PaMsbA and equivalent positions in EcMsbA are shown in orange-colored spheres.

### Supplementary Tables

**Table S1. Equilibrium binding constants for lipids binding to PaMsbA.** $K_D$ was obtained from fitting a sequential lipid binding model to the mole fraction data. Reported are the mean and standard deviation (*n* = 3).

| | $K_{D1}$(μM) | $K_{D2}$(μM) | $K_{D3}$(μM) | $K_{D4}$(μM) | $K_{D5}$(μM) | $K_{D6}$(μM) | $R^{2*}$ | $X^{2*}$ |
|---|---|---|---|---|---|---|---|---|
| **POPA** | 13.6 ± 0.9 | 18.1 ± 1.7 | 23.9 ± 2.9 | 26.8 ± 1.8 | | | 0.98 | 0.07 |
| **POPC** | 48.1 ± 4.2 | 62.3 ± 19.7 | | | | | 0.99 | 0.03 |
| **POPE** | 18.8 ± 2.5 | 33.5 ± 4.6 | 43.0 ± 3.1 | 52.6 ± 7.6 | | | 0.98 | 0.04 |
| **POPG** | 5.9 ± 0.2 | 10.7 ± 0.9 | 16.9 ± 1.2 | 21.0 ± 0.9 | 22.5 ± 1.6 | 22.8 ± 4.9 | 0.97 | 0.06 |
| **TOCDL** | 2.4 ± 0.2 | 4.5 ± 0.6 | 6.3 ± 1.1 | 7.6 ± 0.3 | 8.9 ± 0.7 | 9.8 ± 0.9 | 0.98 | 0.05 |
| **KDL** | 3.0 ± 0.2 | 14.9 ± 3.3 | | | | | 0.99 | 0.04 |

**Table S2. Metal analysis of MsbA samples before and after trapping.** Reported are the mean and standard deviation in units of ng/mL.

| | **Zn** | **Cu** | **Mg** |
|---|---|---|---|
| **EcMsbA** | 0.81 ± 0.13 | 1.93 ± 0.14 | 1.86 ± 0.14 |
| **Trapped EcMsbA** | 0.98 ± 0.04 | 5.49 ± 0.07 | 9.64 ± 0.10 |
| **PaMsbA** | 2.73 ± 0.07 | 1.57 ± 0.01 | 1.91 ± 0.05 |
| **Trapped PaMsbA** | 4.15 ± 0.03 | 1.32 ± 0.03 | 1.13 ± 0.05 |

**Table S3. Equilibrium binding constants for lipids binding to PaMsbA trapped with ADP, $Zn^{2+}$, and vanadate.** $K_D$ was obtained from fitting a sequential lipid binding model to the mole fraction data. Reported are the mean and standard deviation (*n* = 3).

| | $K_{D1}$(μM) | $K_{D2}$(μM) | $K_{D3}$(μM) | $K_{D4}$(μM) | $K_{D5}$(μM) | $R^{2*}$ | $X^{2*}$ |
|---|---|---|---|---|---|---|---|
| **POPA** | 2.5 ± 0.3 | 7.3 ± 0.5 | 10.2 ± 0.7 | 12.7 ± 0.7 | 14.6 ± 0.6 | 0.96 | 0.09 |
| **POPC** | 29.7 ± 4.6 | 35.4 ± 5.6 | 40.8 ± 9.7 | | | 0.98 | 0.08 |
| **POPE** | 6.6 ± 1.1 | 12.6 ± 2.2 | 19.9 ± 2.3 | 26.7 ± 2.4 | 27.1 ± 1.0 | 0.90 | 0.24 |
| **POPG** | 3.9 ± 0.4 | 6.0 ± 0.3 | 8.7 ± 0.1 | 11.9 ± 0.3 | 14.2 ± 0.4 | 0.97 | 0.06 |
| **TOCDL** | 0.4 ± 0.1 | 1.1 ± 0.1 | 1.6 ± 0.2 | 2.8 ± 0.3 | 3.9 ± 0.4 | 0.85 | 0.27 |
| **KDL** | 3.2 ± 0.1 | 14.9 ± 4.3 | | | | 1.00 | 0.01 |

**Table S4. Statistics of cryoEM data collection and processing.**

| | PaMsbA Structures |
|---|---|
| **Microscope** | Titan Krios G4 (Texas A&M University) |
| **Magnification** | 105,000 |
| **Voltage (kV)** | 300 |
| **Spherical aberration (mm)** | 2.7 |
| **Detector** | Gatan K3 |
| **Camera mode** | Super resolution counting |
| **Exposure rate ($e^-$/pixel/s)** | 14.634 |
| **Total exposure ($e^-$/Å$^2$)** | 50.73 |
| **Defocus range (μm)** | -1 to -2.4 |
| **Pixel size (Å)** | 0.832 |
| **Mode of data collection** | Image shift |
| **Energy filter** | 15 eV slit (BioContinuum) |
| **Software for data collection** | EPU |
| **Number of micrographs** | 3,138 |
| **Symmetry imposed** | C2 |
| **Box size (pixel)** | 200 |
| **Initial particle images (no.)** | 1,915,839 |
| **Particle images for 3D (no.)** | 1,334,812 |
| **Final particle images (no.)** | |
| Occluded, outward facing | 506,675 |
| Open, outward-facing | 643,964 |
| **Map resolution, unmasked (Å)** | 4.0 |
| **Map resolution, masked (Å)** | 3.6 |
| **B-factor used for sharpening (Å$^2$)** | 213.7 |
| **EMD (PDB) accession codes** | |
| Occluded, outward-facing | EMD-44444 (9BD6) |
| Open, outward-facing | EMD-44445 (9BD7) |

**Table S5. Statistics of cryoEM model refinement and geometry for PaMsbA structures.**

| Model | Occluded, outward-facing | | Open, outward-facing | |
|---|---|---|---|---|
| Composition (#) | | | | |
| Chains | 4 | | 4 | |
| Atoms | 9036 (Hydrogens: 28) | | 9040 (Hydrogens: 120) | |
| Residues | Protein: 1150 Nucleotide: 2 | | Protein: 1130 Nucleotide: 2 | |
| Water | 0 | | 0 | |
| Ligands | ZN: 2 | | ZN: 2 | |
| | AD9: 2 | | LMT: 2 | |
| | | | AD9: 2 | |
| Bonds (RMSD) | | | | |
| Length (Å) (# > 4σ) | 0.004 (0) | | 0.003 (0) | |
| Angles (°) (# > 4σ) | 0.945 (0) | | 0.480 (0) | |
| MolProbity score | 1.90 | | 1.96 | |
| Clash score | 16.25 | | 15.29 | |
| Ramachandran plot (%) | | | | |
| Outliers | 0.00 | | 0.00 | |
| Allowed | 2.98 | | 1.78 | |
| Favored | 97.02 | | 98.22 | |
| Rama-Z (Ramachandran plot Z-score, RMSD) | | | | |
| whole | (N = 1142) | -1.47 (0.25) | (N = 1122) | 0.75 (0.25) |
| helix | (N = 728) | -0.89 (0.20) | (N = 732) | 1.24 (0.19) |
| sheet | (N = 82) | 1.33 (0.54) | (N = 84) | 0.49 (0.53) |
| loop | (N = 332) | -1.43 (0.32) | (N = 306) | -1.25 (0.32) |
| Rotamer outliers (%) | 1.05 | | 2.25 | |
| Cβ outliers (%) | 0.00 | | 0.00 | |
| Peptide plane (%) | | | | |
| Cis proline/general | 0.0/0.0 | | 0.0/0.0 | |
| Twisted proline/general | 0.0/0.0 | | 0.0/0.0 | |
| CaBLAM outliers (%) | 1.94 | | 0.90 | |
| ADP (B-factors) | | | | |
| Iso/Aniso (#) | 9008/0 | | 8920/0 | |
| min/max/mean | | | | |
| Protein | 19.41/137.39/58.03 | | 9.17/132.17/44.67 | |
| Ligand | 33.34/70.09/41.43 | | 12.60/63.19/38.97 | |
| Data | | | | |
| Box | | | | |
| Lengths (Å) | 85.70, 75.71, 135.62 | | 84.03, 79.04, 134.78 | |
| Angles (°) | 90.00, 90.00, 90.00 | | 90.00, 90.00, 90.00 | |
| Supplied Resolution (Å) | 3.0 | | 2.7 | |
| Resolution Estimates (Å) | Masked | | Masked | |
| d FSC (half maps; 0.143) | 3.0 | | 2.7 | |
| d 99 (full/half1/half2) | 3.2/2.3/2.3 | | 2.9/2.9/2.9 | |
| d model | 3.3 | | 3.0 | |
| d FSC model (0/0.143/0.5) | 2.9/2.9/3.8 | | 2.6/2.7/2.8 | |
| Map min/max/mean | -1.38/2.20/0.03 | | -1.54/2.66/0.03 | |
| Model vs. Data | | | | |
| CC (mask) | 0.86 | | 0.88 | |
| CC (box) | 0.62 | | 0.66 | |
| CC (peaks) | 0.62 | | 0.66 | |
| CC (volume) | 0.83 | | 0.83 | |
| Mean CC for ligands | 0.73 | | 0.74 | |
